# Niche differentiation following whole-genome duplication? The importance of considering the intricate evolutionary history of polyploids when assessing climatic niche evolution

**DOI:** 10.1101/2022.02.16.480351

**Authors:** Nélida Padilla-García, Gabriela Šrámková, Eliška Záveská, Marek Šlenker, Josselin Clo, Vojtěch Zeisek, Magdalena Lučanová, Ieva Rurane, Filip Kolář, Karol Marhold

## Abstract

**Aim:** Although whole genome duplication (WGD) is an important speciation force, we still lack a consensus on the role of niche differentiation in polyploid evolution. In addition, the role of genome doubling *per se* vs. later divergence on polyploid’s niche evolution remains obscure. One reason for this might be that the intraspecific genetic structure of polyploid complexes and interploidy gene flow is often neglected in ecological studies. Here, we aim to investigate to which extent these evolutionary processes impact our inference on niche differentiation of autopolyploids.

**Location:** Europe

**Taxon:** *Arabidopsis arenosa* (Brassicaceae)

**Methods:** Leveraging a total of 352 cytotyped populations of diploid-autotetraploid *A. arenosa*, we examined differences among climatic niches of diploid and tetraploid lineages both globally, and independently for each tetraploid lineage with respect to the niche of its evolutionary closest relative. Then, we tested if there was an effect of additional interploidy introgression from other sympatric but more ancestral diploid lineages of *A. arenosa* on climatic niches of tetraploids.

**Results:** Ecological niche shift of tetraploids is only detected when the assignment of populations to intraspecific genetic lineages is considered. We found different patterns of climatic niche evolution (i.e. niche conservatism, contraction or expansion) in each tetraploid lineage when compared to its evolutionary closest relatives. We observed an effect of interploidy gene flow in patterns of climatic niche evolution of tetraploid ruderal plants of *A. arenosa*.

**Main conclusions:** The niche shift of tetraploids in *A. arenosa* is not driven by WGD per se but rather reflects dynamic post-WGD evolution in the species, involving tetraploid migration out of their ancestral area and interploidy introgression with other diploid lineages. Our study supports that evolutionary processes following WGD - which usually remain undetected by studies neglecting evolutionary history of polyploids - may play a key role in the adaptation of polyploids to challenging environments.

## INTRODUCTION

Polyploidy is a leading evolutionary force driving speciation and diversification of all plant lineages (Otto & Whitton, 2000; Wendel, 2000; Soltis, Marchant, Van de Peer & Soltis, 2015). All flowering plants have experienced one or more episodes of whole-genome duplication (WGD) during their evolutionary history (Jiao et al., 2011; Wendel, 2015) and it has been estimated that up to 15% of all speciation events in angiosperms are associated with a ploidy increase (Wood et al., 2009). Polyploid speciation has been considered a mechanism of sympatric speciation (Coyne & Orr, 2004) given the potential reproductive isolation that newly formed polyploids can experience due to strong postzygotic barriers among cytotypes (e.g. differences in the number of sets of chromosomes). However, incomplete postzygotic isolation among cytotypes is common in plants, thus promoting interploidy reproduction (Sutherland & Galloway, 2017). Accordingly, polyploid speciation is not an “instantaneous” process and prezygotic reproductive barriers such as differences in ecological niches between polyploids and closest lower-ploidy progenitors can also play a key role in assortative mating and thus, polyploid speciation (Levin, 2004; Husband, Baldwin & Sabara, 2016). Novel phenotypic, physiological and genetic combinations associated with polyploidy potentially could drive up polyploids to expand to new ecological niches that would have remained unavailable to their diploid progenitors (Madlung, 2013). In line with this, niche ecological divergence with respect to progenitors may be more prone to occur in allopolyploids (i.e. formed by genome duplication after hybridization of two different parental genomes more or less divergent) than autopolyploids (i.e. formed within the same species or genetic lineage) because a high heterozygosity could allow allopolyploids to colonize new habitats. Many empirical studies have addressed niche evolution after WGD, with some of them supporting niche divergence between diploids and polyploids (Theodoridis, Randin, Broennimann, Patsiou & Conti, 2013; Thompson, Husband & Maherali, 2014; Visger et al., 2016; Muñoz-Pajares et al., 2018; Decanter, Colling, Elyinger, Heiðmarsson & Matthies, 2020), while others do not (Godsoe, Larson, Glennon & Segraves, 2013; Glennon, Ritchie & Segraves, 2014; Čertner, Kolář, Schönswetter & Frajman, 2015; Visser & Molofsky, 2015; Castro et al., 2019; 2020a). Yet, there is not a consistent or universal pattern of the role of niche differentiation in polyploid establishment and evolution. This incongruence between studies has been explained by methodological issues as an inappropriate resolution of environmental variables (Kirchheimer et al., 2016), because niche differentiation might be occurring at a different spatial scale (Čertner, Kúr, Kolář, & Suda, 2019) and/or simply by different evolutionary histories of polyploid cytotype between studied species - an aspect that has been, however, often neglected in purely ecological studies. The presence or absence of niche evolution in polyploids strongly depends on species’ history: e.g. polyploid’s age, multiple origins or the number of ploidy levels, among others (see Duchoslav et al., 2020). There can also be a phylogenetic signal in environmental traits (Burns & Strauss, 2011), thus, ecological niches of polyploids can be potentially affected by ancestral niches of progenitors. Consequently, patterns of ecological differentiation could be misunderstood if niches of the polyploids are not compared to their closest lower-ploidy progenitors. Unfortunately, the intraspecific genetic structure of polyploid complexes is often unknown due to limitations associated with challenges surrounding population genetic data analysis of polyploids (Rothfels, 2021; Rojas-Andrés et al., 2020), and if known, it is rarely taken into account when evaluating polyploid niche evolution (see López-Jurado, Materos-Naranjo & Balao, 2019 as an exception). Niches of allopolyploids are predicted by the sum of the niche of their progenitors (Parisod & Broennimann, 2016). This is not the case of strict autopolyploids, which usually conserve the climatic niche of their unique progenitor species. Nevertheless, hybridization and interploidy admixture between geographically close diploid and tetraploid populations are common in many plant species (Aagaard, Såstad, Greilhuber & Moen, 2005; Stahlberg, 2007; Koutecký, Baďurová, Štech, Košnar & Karásek, 2011; Monnahan et al., 2019; Šmíd, Douda, Krak & Mandák, 2020). Thus, it might also occur that autopolyploids come into secondary contact with diploid ancestors that have diverged before the WGD event. This genetic exchange may result in adaptive introgression that can also influence patterns of climatic niche evolution (Schmickl & Yant, 2021). Nevertheless, few studies have assessed whether introgression promotes ecological niche evolution of polyploids, most of them being focused on strict allopolyploids (Arrigo et al., 2016; Blaine Marchant, Soltis & Soltis, 2016; Manzoor, Griffiths, Obiakara, Esparza-Estrada & Lukac, 2020). The effect of interploidy introgression in the ecological niche of autopolyploids remained unexplored.

*Arabidopsis arenosa* (Brassicaceae) is a diploid-autotetraploid species with a well-described evolutionary history of its lineages (see Fig. 1), that has recently become an interesting system not only to study polyploid evolution (Monnahan et al., 2019; Morgan et al., 2021a; 2021b; Bohutínská et al., 2021a), but also adaptation to extreme conditions (Konečná et al., 2021, Bohutínská et al., 2021b, Knotek et al., 2020, Wos et al., 2021). Multiple studies have addressed the role of niche differentiation in the autopolyploid evolution of this system, however, reaching strikingly inconsistent outcomes. Ecological niche modeling was used in a study conducted by Molina-Henao & Hopkins (2019), which concluded niche expansion but not divergence of tetraploids of *A. arenosa*. These results contrast with other studies in which an absence of ecological niche differentiation was described at both the landscape (Kolář et al., 2016b) and intra-population scales (Wos, Bohutínská, Nosková, Mandáková & Kolář, 2019), and in three contact zones independently (Morgan et al., 2020). Unfortunately, neither of these studies did cover the whole distribution range of the species (Wos et al., 2019; Morgan et al., 2020) and they did not integrate the intricate evolutionary history of this species involving the intraspecific genetic sub-structure of each cytotype and interploidy gene flow (Kolář et al., 2016b; Molina-Henao & Hopkins 2019). Autotetraploid cytotype of *A. arenosa* originated only once in the Western Carpathians 20,000 to 31,000 generations ago, where it still coexists with its diploid progenitor until now, but also from where it spread through most of Europe from Romania in the south to Belgium in the west and Scandinavia in the north (Arnold, Kim & Bomblies., 2015; Monnahan et al., 2019, summarized in Fig. 1). During this expansion, tetraploids also encountered other earlier diverged diploid lineages of *A. arenosa* and got introgressed by them in at least two contact zones, in SE Carpathians and the Baltic coast (Monnahan et al., 2019). An intriguing question about to which extent such intricate evolutionary history has been translated to niche divergence, however, remained unanswered.

**Figure 1.**
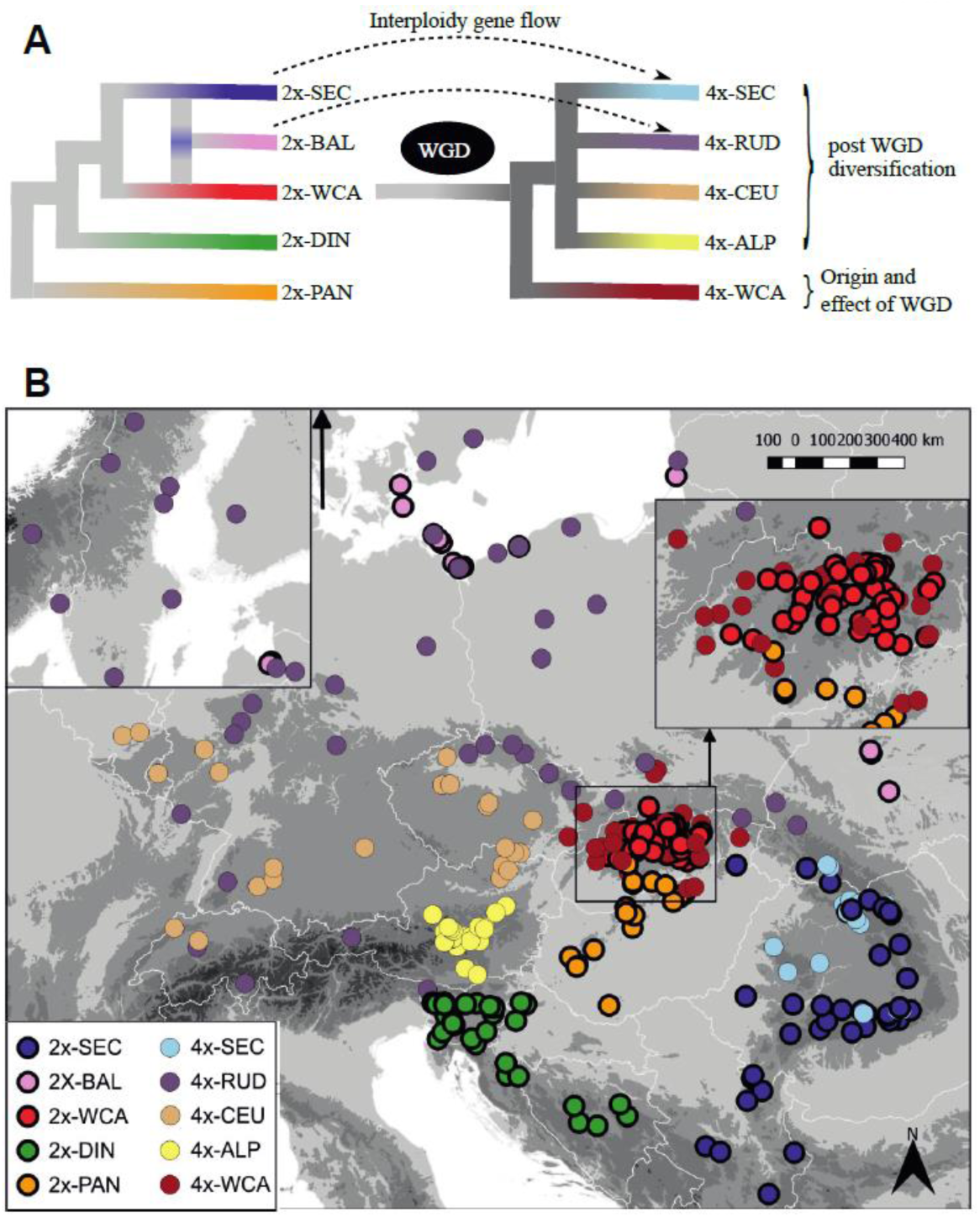
**A** Summary of evolutionary history of diploid and autotetraploid lineages within *A. arenosa*. Based on our previous knowledge, we examined the impact of the intraspecific genetic structure and the interploidy introgression when testing for autopolyploid climatic niche differentiation. A single whole-genome duplication event occurred within the species is indicated by “WGD”. Interploidy gene flow events are indicated by dashed lines; **B** Geographic distribution of populations from diploid and autotetraploid lineages of *A. arenosa* included in our niche quantification and comparison analyses is shown.

In the present study, we aim to examine the impact of the intraspecific genetic structure and interploidy introgression when testing for autopolyploid niche evolution. More specifically, we test the following hypothesis: (i) whether niche shift of polyploids is detectable only when polyploid lineages are compared with their corresponding diploid ancestor(s), not globally; and (ii) whether interploidy introgression events promote ecological divergence and/or expansion of polyploids. First, we used genome-wide single nucleotide polymorphism (SNP) genotyping to investigate the genetic structure within *A. arenosa* tetraploids across the landscape including for the first time samples from the whole geographic distribution area of the tetraploid cytotype. Second, we compared the environmental niche occupied by diploid and tetraploid lineages of *A. arenosa*, both globally, and independently for each tetraploid lineage with respect to the niche of the evolutionary closest diploid lineage, and among them. Third, we investigated the effect of interploidy gene flow in polyploid niche shift by comparing niches of tetraploid Ruderal and SE Carpathian lineages to the niches of its respective sympatric locally-adapted diploid-related lineages.

## MATERIAL AND METHODS

### Genomic data

#### Dataset and library preparation

The cytogeographical patterns in *A. arenosa* have been well documented (Kolář et al., 2016b) and the genetic structure of diploid populations across their entire distributional range is well known (Kolář et al., 2016a). However, in the case of tetraploids, the precise identification of genetic lineages still remains unclear mainly because the whole distribution range of tetraploids of *A. arenosa* has not been fully covered in previous studies (Arnold et al., 2015), or the number of sampled populations was limited (Monnahan et al., 2019). In order to identify range-wide tetraploid genetic structure, we generated a dataset of genome-wide single-nucleotide polymorphisms (SNPs) for a total of 276 tetraploid individuals from 125 populations (1-4 indiv/pop) using double-digest RADseq (as described in Wos et al., 2019). On top of that, available published WGS data (Monnahan et al., 2019) from 62 tetraploid populations (155 individuals) were also integrated to generate a more robust dataset, consisting of a total of 431 individuals from 187 populations.

#### Raw data processing, variant calling and filtration

Raw reads were demultiplexed using FASTX toolkit 0.0.14 and quality trimmed (> 20 Phred quality score) in Trimmomatic 0.36. Mapping on *Arabidopsis lyrata* reference genome v. 1.0.25 (Hu et al., 2011) was performed using BWA v. 0.7.3a and the resulting BAM was processed with Picard Tools v. 2.22.1. The Genome Analysis Toolkit v. 3.8 (GATK, McKenna et al., 2010) was used following the best practice recommendations (https://gatk.broadinstitute.org/hc/en-us). Variant calling was performed for each individual using the HaplotypeCaller module, setting the ploidy=4 option. Then, we aggregated variants and performed genotyping across all individuals using GenotypeGVCFs.

The coordinates of the identified RAD loci were used to retrieve the corresponding SNPs from a previous set of genome resequencing data (Monnahan et al., 2019) that was mapped to the same reference genome. In this way, we extracted the SNP data from the same sites for an additional 155 individuals using GATK. Both VCF files were merged and the final VCF contained 431 tetraploid individuals from a total of 187 populations. Only biallelic sites that passed the filter parameters indicated by GATK best practices (https://gatk.broadinstitute.org/hc/en-us/articles/360035890471-Hard-filtering-germline-short-variants) were considered: ‘QD < 2.0’, ‘FS > 60.0’, ‘MQ < 40.0’, ‘MQRankSum < -12.5’, ‘ReadPosRankSum < -8.0’, ‘SOR > 3.0’. Additionally, for subsequent analyses, only variants that were present in at least 80% of individuals at a minimum sequencing depth of 8x were used, reaching a final dataset of 179,698 SNPs. Scripts used for processing the data are available at https://github.com/V-Z/RAD-Seq-scripts.

#### Identification of genetic lineages

Population genetic structure was inferred using Bayesian clustering analyses in STRUCTURE v. 2.3.2 (Pritchard, Stephens & Donnelly, 2000), which allowed us to take into consideration autotetraploid genotypes. Before running STRUCTURE, the dataset was pruned to avoid linkage among SNPs. Taking into account the average length of RADseq fragments (350 bp), we randomly selected one SNP per each 1000-bp window to avoid linkage disequilibrium. We discarded those SNPs showing a lower minor allele frequency of 0.05 and a higher minor frequency of 0.95 to remove uninformative singletons and errors in the dataset. Python scripts used for pruning and formatting the input data for STRUCTURE are available at https://github.com/MarekSlenker/vcf_prune. In STRUCTURE, we run ten replicates per each value of *K* between 1 and 10 applying a burn-in of 10^4^ iterations followed by 10^5^ MCMC iterations. Convergence among different replicates per each *K* value was evaluated in R using a script (https://github.com/MarekSlenker/structureSum) that ran modified functions previously coded by Ehrich (2006). The results for every value of *K* were visualized using CLUMPAK (Kopelman, Mayzel, Jakobsson, Rosenberg & Mayrose, 2015). The optimal value of groups in our dataset was identified according to several criteria (i.e. *K* = 5 is the highest value of *K* showing a positive of the results among the ten replicates; Fig. S1). Each population was assigned to the cluster for which the highest proportion of membership was observed. We further investigated the population structure inferenced by STRUCTURE running principal component analysis (PCA). Based on putatively neutral four-fold degenerate (4dg) SNPs and using ADEGENET package in R (Jombart, 2008) we summarized the neutral genetic variability among the identified lineages within our tetraploid samples. Additionally, we calculated pairwise Nei’s genetic distances (Nei, 1972) among the same lineages to quantify the genetic differentiation between them using StAMPP R package (Pembleton, Cogan & Forster, 2013).

### Niche comparison analyses

#### Occurrence data and climatic variables

For the climatic niche comparison analyses, we used both diploid and tetraploid occurrences of *A. arenosa* with known affiliations to the genetic lineages (Supplementary Table 1). We collected leaf material from diploid and tetraploid populations of *A. arenosa* in 2011–2020 across their entire European distribution range and we checked the ploidy level of each individual using flow cytometry as described in Kolář et al. (2015). Geographical coordinates were obtained from GPS during field surveys. The lineage assignment to tetraploid populations was done according to the Bayesian clustering of the SNP data obtained in this study (see details above and Supplementary Table 2). To avoid potential bias caused by admixture between different tetraploid lineages, we excluded equivocal populations showing less than 50% membership to one single cluster from the niche comparison analyses (in total 21 populations, see Supplementary Table 2). In case of diploids, the assignment was based on previous Bayesian clustering of a set of populations covering the entire range of the diploid cytotype (Kolář et al., 2016a) and the geographical location of additionally sampled populations. A total of 352 localities including 2*x* and 4*x* cytotypes of *A. arenosa* assigned to intraspecific genetic lineages were analyzed (see Supplementary Table 1). In order to avoid unequal representation caused by biased sampling in different areas, the obtained sampling points were filtered and those that were closer than 10-km distance were removed. The number of occurrences per cytotype and lineage is indicated in Supplementary Table 3. Environmental data related to temperature (BIO1-BIO11 variables) and precipitation (BIO12-BIO19 variables) were extracted for all occurrence points from WorldClim at 30-s (ca. 1 km) resolution (Hijmans, Cameron, Parra, Jones & Jarvis, 2005).

#### Niche quantification and comparison

Several climatic niche comparisons were performed: i) between diploids and tetraploids of *A. arenosa* globally, without considering the assignment to intraspecific genetic lineages; ii) independently between each tetraploid lineage and its evolutionary closest diploid relative; iii) among tetraploid lineages that diverged after the WGD event; and iv) between tetraploid and locally sympatric diploid lineages. Then, we tested if there was an effect of other sympatric but more ancestral diploid lineages of *A. arenosa* on climatic niches of tetraploids. Quantification and comparison of climatic niches were performed using a statistical framework developed by Broennimann et al. (2012). It applies a kernel density function to calculate the smoothed density of occurrences and environmental values along the first two axes of a multivariate analysis (PCA-env). This method ensures that the niche overlap is independent of the resolution of the grid. We considered the first two axes of the PCA calibrated on the environmental space of the study area, which was divided into a grid of 100 × 100 cells with each cell corresponding to a unique set of environmental conditions. The environmental space was produced by extracting the same climatic values for 10,000 occurrence points randomly sampled from 100-km buffer zones around the occurrences of diploid and tetraploid *A. arenosa* records. This common environmental space, which theoretically corresponds to the potential habitat of the species, was used for all pairs of comparisons. Occurrence density grids had a resolution of 100 and a species density threshold of zero. Niche overlap, equivalence and similarity tests implemented in the R package “ecospat” (Di Cola et al., 2017) were performed to compare the divergence between lineages. Niche overlap calculation is based on Schoener’s *D* metric (Schoener, 1968) that ranges from 0 (no overlap) to 1 (complete overlap). To evaluate the significance of niche overlap (α = 0.05), we performed an equivalency test (Warren, Glor & Turelli, 2008), which uses random bootstrap resampling of presence occurrence points of both lineages to calculate if a null distribution of *D* and the observed *D* are significantly different (p < 0.05; niches are not statistically equivalent) or not (niches are equivalent). The significance of niche overlap was also evaluated by similarity tests, which use bootstrap resampling to assess whether the niche of one lineage predicts the other better than would be expected by chance (α = 0.05). If observed *D* is greater than the null distribution, niches are more similar than expected. Values lower than the null distribution indicate that niches are not similar and non-significant values mean a lack of power of the test to detect differences or similarities. However, simply testing if niches of diploid and tetraploid cytotypes are equivalent or different does neither fully account for dynamics of niche evolution, nor reflects complex reticulated evolutionary history of the species. In order to understand alternative processes driving niche evolution in *A. arenosa* we have quantified niche dynamics per each genetic lineage, using niche unfilling (U), stability (S) and expansion (E) indices (Guisan, Petitpierre,Broennimann, Daehler & Kueffer, 2014) and calculating niche optimum and breadth along the axes of temperature and precipitation variation. Niche optimum and breadth of each lineage was calculated following the procedure described in Theodoridis et al. (2013) and Kirchheimer et al. (2016). We randomly sampled 100 cells of the gridded space of each lineage with the probability of selection weighted by the density of the species occurrences. We calculated the niche optimum and breadth, calculating the median and the standard deviation of the scores along the two PCA axes, respectively. This re-sampling was repeated 1000 times and differences in the distribution of optimum and breadth values were compared using Welch’s t-tests in R. The results were visualized using boxplots for each PCA axis. In order to test for an effect of the studied lineages on the niches’ optimum and breadth for each PCA axis, we first performed an ANOVA test, and then we performed a Tukey HSD test to perform all the possible pairs-comparisons.

## RESULTS

### Genetic structure within *A. arenosa* tetraploids

We have obtained a total of 179,698 filtered SNPs for the 431 tetraploid individuals of *A*.*arenosa* included in our study. The average depth is 11× and our dataset comprises 13.1% of missing data over populations and sites. Bayesian clustering analyses support the existence of five distinct genetic clusters among tetraploids of *A. arenosa*. These clusters are geographically separated (Fig. 2). Two of them (dark red and light blue) are restricted to populations located in W and SE Carpathians, respectively. The majority of populations from the Western part of Central Europe (further referred to as C Europe) are included in a third cluster (light orange). Only populations from the Eastern Alps form a separate fourth cluster (light yellow). Finally, some populations sampled in human-made ruderal stands (mainly railways tracks and roadsides) in the Alps, Germany and the Czech Republic are clustered together with all populations located in Northern Europe and the Baltic Sea coast (purple).

**Figure 2.**
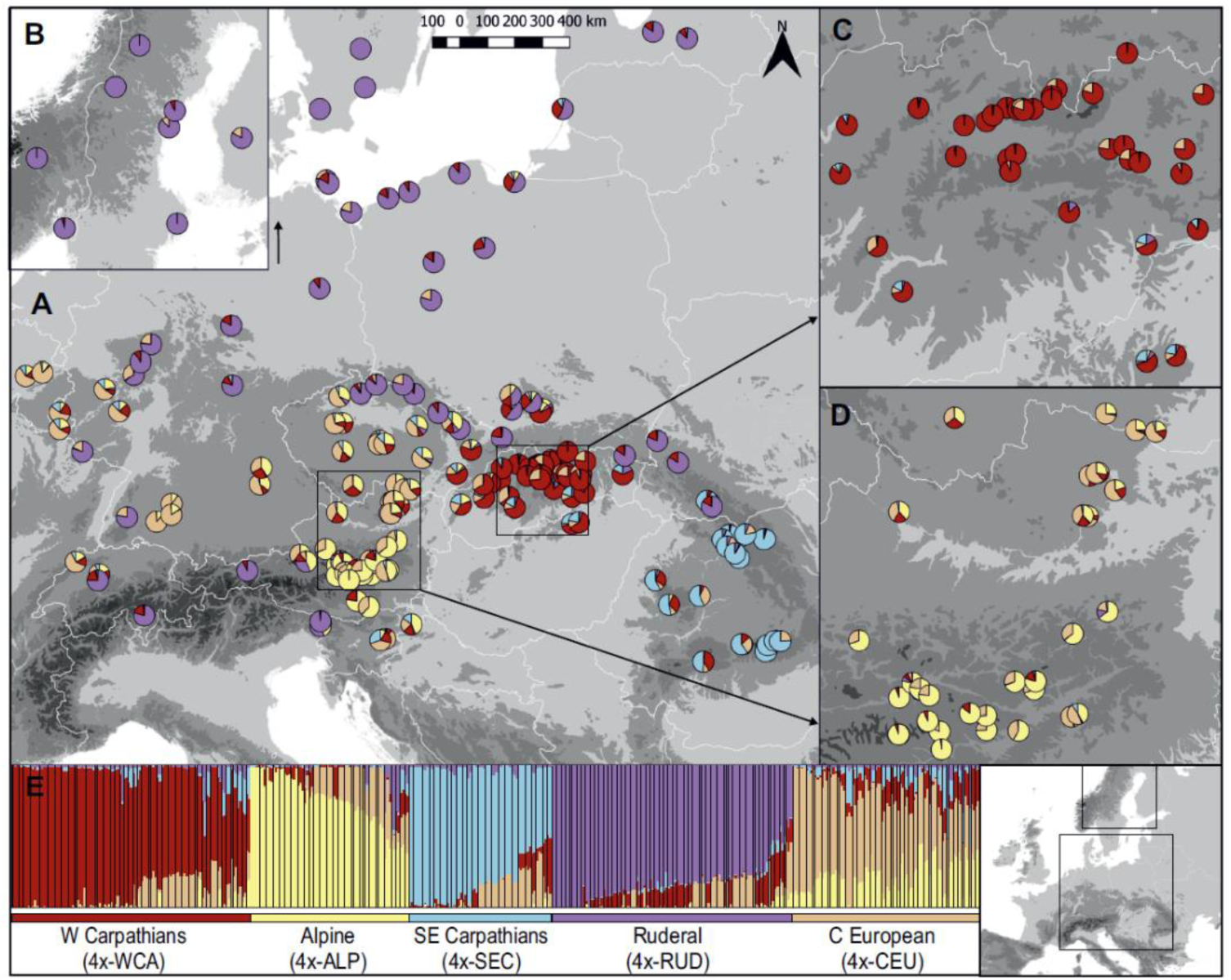
Rangewide genetic differentiation of tetraploid *A. arenosa*. **A** Geographic distribution of populations used in the genomic analyses (color pie charts reflecting the proportional assignment to particular clusters identified by STRUCTURE and representing tetraploid lineages within *A. arenosa*: W Carpathians (dark red), Alpine (light yellow), SE Carpathians (light blue), Ruderal (purple), C European (light orange); **B** Detail of Scandinavia; **C** Detail of the Western Carpathian region; **D** Detail of the Austrian region; **E** Individual assignment into each of the five genetic clusters revealed by STRUCTURE.

Principal Component Analysis based on genetic data confirms the genetic differentiation of *A. arenosa* tetraploids into the five lineages and further indicates the level of differentiation among them. Ruderal populations (4x-RUD) are separated from the others along the first axis, while populations located in SE Carpathians (4x-SEC) are differentiated from the rest along the second axis (Supplementary Figure 2a). Alpine (4x-ALP), W Carpathians (4x-WCA) and C European (4x-CEU) populations cluster together in the PCA of the complete dataset but get clearly distinct in a separate analysis excluding the populations identified within the 4x-RUD and 4x-SEC lineages (Supplementary Figure 2b). Nei’s genetic distances calculated among lineages reflect the fact that the genetic divergence among them is generally low (Supplementary Figure 2c).

### Niche quantification and comparison between diploids and tetraploids globally

The variation explained by the two first axes of the PCA-env is 41.5% and 30.4% respectively, which means a 71.9% of the total variance. Environmental variables related to precipitation (BIO12-BIO19) are highly correlated to PC1, while temperature-related variables (BIO1-BIO11) are mainly correlated to PC2 (Fig. 3a). The contribution of each variable to the two first axes of the PCA is summarized in Supplementary Table 4. Annual precipitation (BIO12), precipitation of the driest quarter and month (BIO17 and BIO14) together with precipitation of the coldest quarter (BIO19) show the highest percentage of correlation to PC1. Mean temperature of the coldest quarter (BIO11) and minimum temperature of the coldest month (BIO6) are the most correlated variables to PC2.

**Figure 3.**
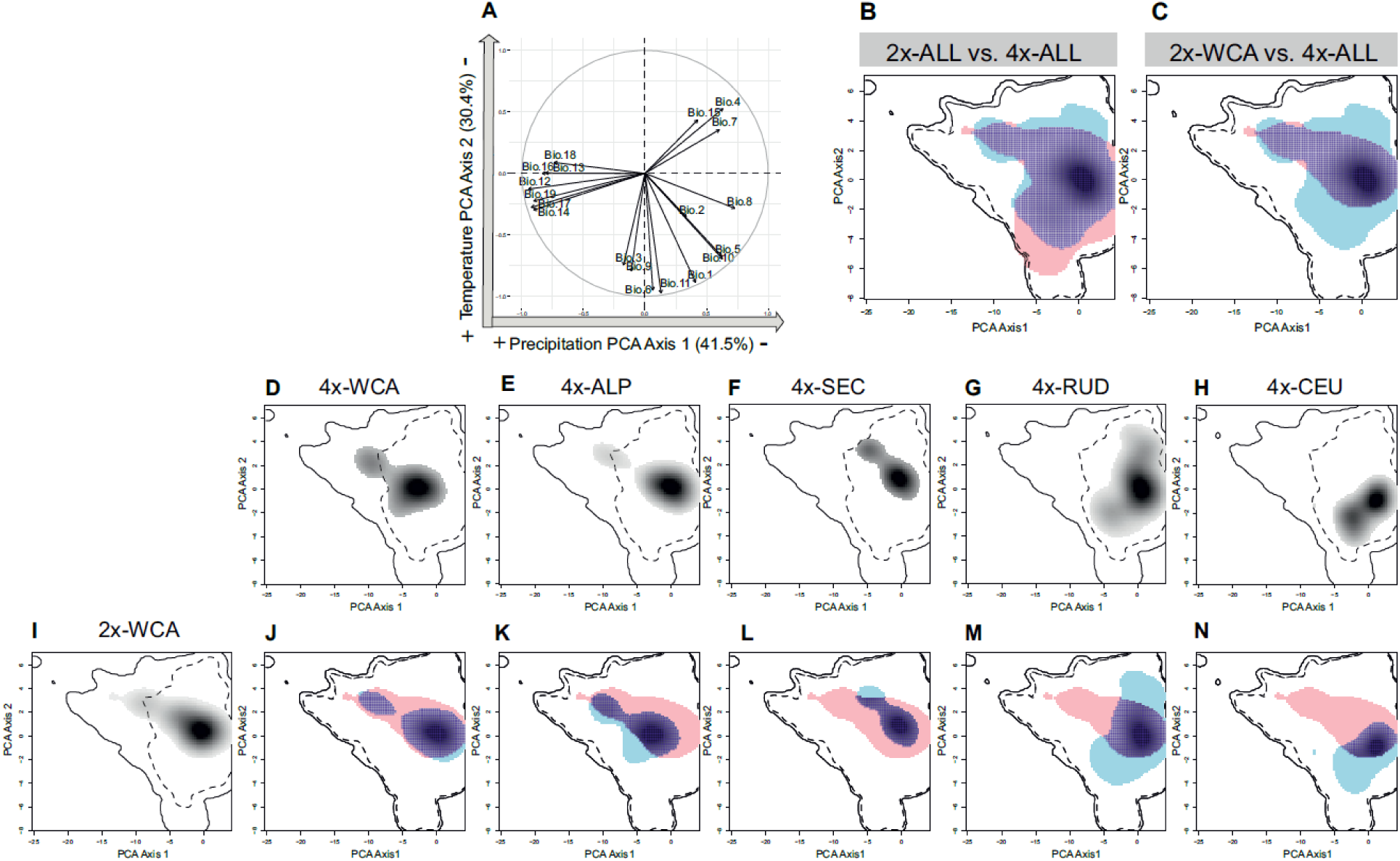
Climatic niche dynamics of diploid and tetraploid *A. arenosa* lineages. The fraction of the diploid niche that remains unfilled by tetraploids represents tetraploid niche contraction (pink), whereas climatic space occupied by the tetraploid and not by the diploids indicates niche expansion (blue). The fraction of the tetraploid climatic niche that overlaps with diploids indicates niche stability (dark blue). **A** Correlation circle of 19 environmental variables along the first two axes of PCA-env. The percentage of variation and the main environmental gradient explained by each axis is indicated; **B** Niche of all *A. arenosa* diploids compared to the niche of all *A. arenosa* tetraploids without considering their assignment to different lineages; **C** Niche of ancestral diploid lineage (2x-WCA) compared to the niche of all *A. arenosa* tetraploids; **D-I** Climatic niche reconstruction of each *A. arenosa* tetraploid lineage and the ancestral diploid W Carpathian lineage. Occurrence density grids are represented by a black-to-white downward gradient along the two first axes of PCA-env; **J-N** Comparisons of climatic niches of each tetraploid lineage to their ancestral diploid lineage.

When the niche of diploids and tetraploids of *A. arenosa* is compared globally, i.e. ignoring the assignment to intraspecific genetic lineages, the niche equivalency test does not have enough power to detect niche differentiation, while the similarity test indicates that niches are more similar than expected by chance independent of the direction of the test (Table 1). These results indicate niche conservatism between diploids and tetraploids when they are globally compared (Fig. 3b). Accordingly, tetraploids show a relative high niche overlap with diploids (50.2 %), very high stability index value (S = 0.947) and almost zero expansion and niche unfilling (E = 0.053 and U = 0.079, respectively).

**Table 1.** Niche overlap percentage values, results for equivalency and similarity tests and niche dynamic indices (expansion, stability and unfilling) are shown for pairwise comparisons (lineage 1 vs. lineage 2).

| *Arabidopsis arenosa* |  | % Niche overlap | Niche equivalency | Niche similarity |  | Indices of niche change |  |  |
| --- | --- | --- | --- | --- | --- | --- | --- | --- |
| 1 | 2 |  |  | 1 → 2 | 2 → 1 | Expansion (E) | Stability (S) | Unfilling (U) |
| 2x-ALL | 4x-ALL | 50.2 | ns (EQUIVALENT) | More similar * | More similar * | 0.053 | 0.947 | 0.079 |
| 2x-WCA | 4x-ALL | 62.8 | ns (EQUIVALENT) | More similar * | More similar * | 0.252 | 0.748 | 0.006 |
| 2x-WCA | 4x-WCA | 64.9 | ns (EQUIVALENT) | More similar * | More similar * | 0.017 | 0.983 | 0.127 |
| 2x-WCA | 4x-ALP | 52.1 | ns (EQUIVALENT) | ns | ns | 0.215 | 0.785 | 0.350 |
| 2x-WCA | 4x-SEC | 24.6 | ns (EQUIVALENT) | ns | ns | 0.097 | 0.903 | 0.400 |
| 2x-WCA | 4x-RUD | 28.1 | ns (EQUIVALENT) | ns | ns | 0.336 | 0.664 | 0.207 |
| 2x-WCA | 4x-CEU | 7.7 | ns (EQUIVALENT) | ns | ns | 0.479 | 0.521 | 0.607 |
| 4x-WCA | 4x-ALP | 46.8 | ns (EQUIVALENT) | ns | ns | 0.329 | 0.672 | 0.347 |
| 4x-WCA | 4x-CEU | 14.4 | ns (EQUIVALENT) | ns | ns | 0.447 | 0.553 | 0.438 |
| 4x-WCA | 4x-RUD | 32.4 | ns (EQUIVALENT) | ns | ns | 0.364 | 0.636 | 0.098 |
| 4x-WCA | 4x-SEC | 21.6 | ns (EQUIVALENT) | ns | ns | 0.258 | 0.742 | 0.390 |

\* The ecological niches are significantly ( $P < 0.05$ ) more similar than expected

### Niche quantification and comparison between related diploid and tetraploid intraspecific lineages

To integrate niche data with evolutionary history (Fig. 1), we firstly reconstructed niche evolution reflecting the single evolutionary origin of all *A. arenosa* tetraploid lineages from the ancestral diploid lineage (2x-WCA). Niche conservatism is observed when comparing all *A. arenosa* tetraploid lineages to the 2x-WCA lineage, but excluding other diploid lineages from the analysis that did not contribute to the origin of autotetraploids (Fig. 3c). In this case, the proportion of niche overlap of tetraploids with the 2x-WCA diploid lineage is 62.8% (Table 1). The stability index value is high (S = 0.748) while the expansion and unfilling values are E = 0.252 and U = 0.006. Interestingly, when each tetraploid lineage was compared independently to the ancestral 2x-WCA lineage, the resulting patterns are very different from the previous global comparison and from each other (Fig. 3). Specifically in the W Carpathians, niches of diploids and tetraploids are more similar than expected by chance, independent of the direction of the test (Table 1). Furthermore, a high niche overlap is observed between 2x and 4x W Carpathians lineages (64.9%) with high stability and very low values for expansion and niche unfilling indices (S = 0.983, E = 0.017 and U = 0.127). These results together suggest that in the W Carpathians, diploid and tetraploid lineages show similar ecological niches. In contrast, similarity tests were non-significant (Table 1) when comparing other tetraploid lineages to the ancestral diploid (2x-WCA). A high overlap was found between niches corresponding to 4x-ALP and 2x-WCA (52.1%), also showing a high stability value (S = 0.785) and intermediate values of niche expansion (E = 0.215) and unfilling (U = 0.350). In the case of 4x-SEC lineage, niche contraction compared to the 2x-WCA lineage is indicated by an intermediate value of unfilling index (U = 0.400) and a high stability index (S = 0.903) but almost zero expansion (E = 0.097). In this case, the percentage of niche overlap is 24.6% (Table 1). The 4x-RUD lineage has experienced niche expansion compared to the 2x-WCA lineage (S = 0.664 and E = 0.336), whose niche was partially filled by the Ruderal one (U = 0.207). These results indicate that niche expansion of the 4x-RUD has occurred towards areas of more extreme temperatures while it has not filled the “high-alpine” niche portion occupied by its 2x-WCA ancestor lineage, which is characterized by high precipitation values. The niche occupied by the 4x-CEU lineage has also significantly expanded (S = 0.521 and E = 0.479), but in this case tetraploids almost did not fill the niche occupied by the 2x-WCA lineage (U = 0.607).

Niche optimum (calculated as the median of the scores along each axis of PCA-env) reflects a higher optimal precipitation value (PC1) for 2x-WCA when compared to all tetraploid lineages, with the exception of the 4x-ALP (Fig. 4). Optimal temperature values (PC2) are higher for tetraploids than for 2x-WCA except for the 4x-SEC lineage. Niche breadth in terms of precipitation (PC1) is lower for all tetraploid lineages compared to the 2x-WCA lineage (Fig. 4). In terms of temperature (PC2), niche breadth is not significantly different among 2x-WCA, 4x-WCA and 4x-CEU lineages. Temperature breadth is higher for 2x-WCA than for the 4x-ALP and 4x-SEC lineages but lower than for the 4x-RUD one.

**Figure 4.**
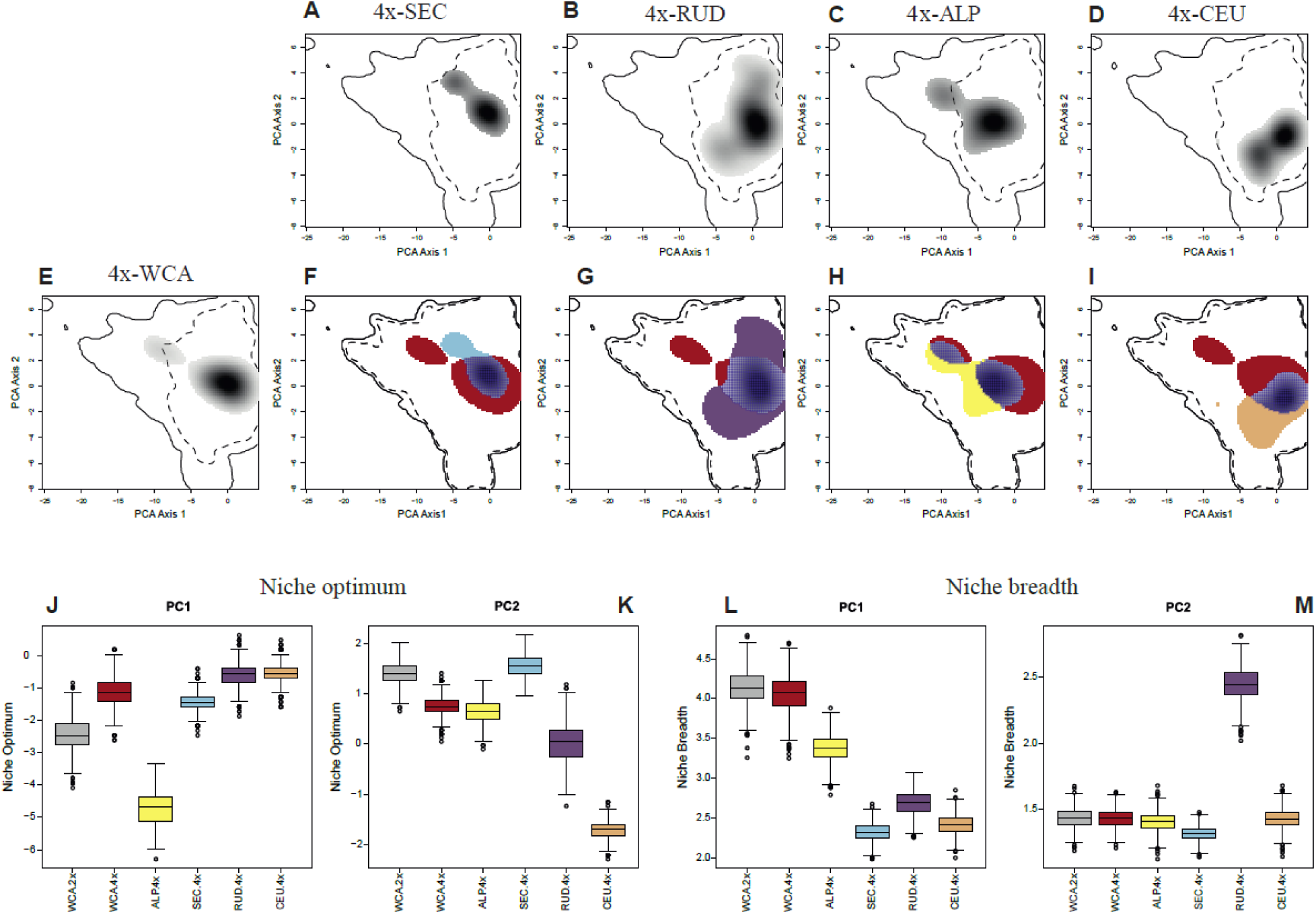
**A-E** Climatic niche reconstruction for each tetraploid lineage of *A. arenosa*. Occurrence density grids are represented by a black-to-white downward gradient along the two first axes of PCA-env. **F-I** Niche comparisons between each tetraploid lineage and the ancestral W Carpathian tetraploid. The fraction of the W Carpathian tetraploid niche that remains unfilled is represented in dark red, whereas niche expansion of each tetraploid lineage is colored according to Fig. 2. The niche overlap is indicated in dark blue. **J-M** Niche optimum and breadth for W Carpathian diploid and each of the tetraploid *A. arenosa* lineages along the PC1 and PC2 axes.

### Niche quantification and comparison among tetraploid lineages

Furthermore, we inferred niche evolution during tetraploid expansion. To do so, we compared the niche of each tetraploid lineage to the niche of tetraploids from the W Carpathians (4x-WCA) which occupy the presumed area of origin of tetraploid *A. arenosa* cytotype and are thus closest to the “ancestral polyploid” niche (Fig. 1). Similarity tests were not significant for any pair-wise comparison (Table 1). Percentages of niche overlap varied from 14.4 % (4x-CEU) to 46.76 % (4x-ALP). All tetraploid lineages showed some degree of niche expansion relative to the 4x-WCA lineage occupying the ancestral area (Fig. 4), with values of E indices ranging between 0.258 (4x-SEC) and 0.447 (4x-CEU). The 4x-RUD lineage almost entirely filled the niche of 4x-WCA (U = 0.098) while other lineages show values of unfilling indices that vary between 0.347 (4x-ALP) and 0.438 (4x-CEU). The optimal value of precipitation (PC1) is shown to be highest for 4x-ALP and lowest for 4x-RUD and 4x-CEU lineages, while in terms of temperature (PC2) is the 4x-SEC which showed the coldest optimal value. Specifically, 4x-RUD and 4x-CEU lineages show the highest optimal values of temperature (Fig. 4). Niche breadth indicates that 4x-WCA occupies the broadest gradient of precipitation compared to all other tetraploid lineages, while the temperature breadth is much higher for the 4x-RUD lineage than from other tetraploid lineages (Fig. 4).

### Niche quantification and comparison between tetraploid and locally sympatric diploid lineages

Last, we investigated niche evolution when taking into account the influence of strong inter-ploidy introgression from more divergent diploid lineages of *A. arenosa* which co-occur with particular tetraploid lineages within two contact zones (Fig. 1): SE Carpathians (4x-SEC lineage introgressed by 2x-SEC) and Baltic coast (4x-RUD lineage introgressed by 2x-BAL). When we compared the observed niche of the 4x-SEC lineage to the combined niche of both diploid lineages that served as source gene pools (2x-WCA and 2x-SEC), we do not observe substantive changes in niche dynamics index values with respect to the previous comparison including 4x-SEC lineage and only the ancestral diploid 2x-WCA lineage (Fig. 3l, Fig. 5a). It is shown that tetraploids have barely expanded their niche (S = 0.923 and E = 0.077) and that a considerable proportion of the niche of the diploid lineages has not been filled by 4x-SEC (U = 0.417). Thus, niche contraction with respect to both diploid lineages has occurred in the 4x-SEC lineage towards warmer and more humid areas.

**Figure 5.**
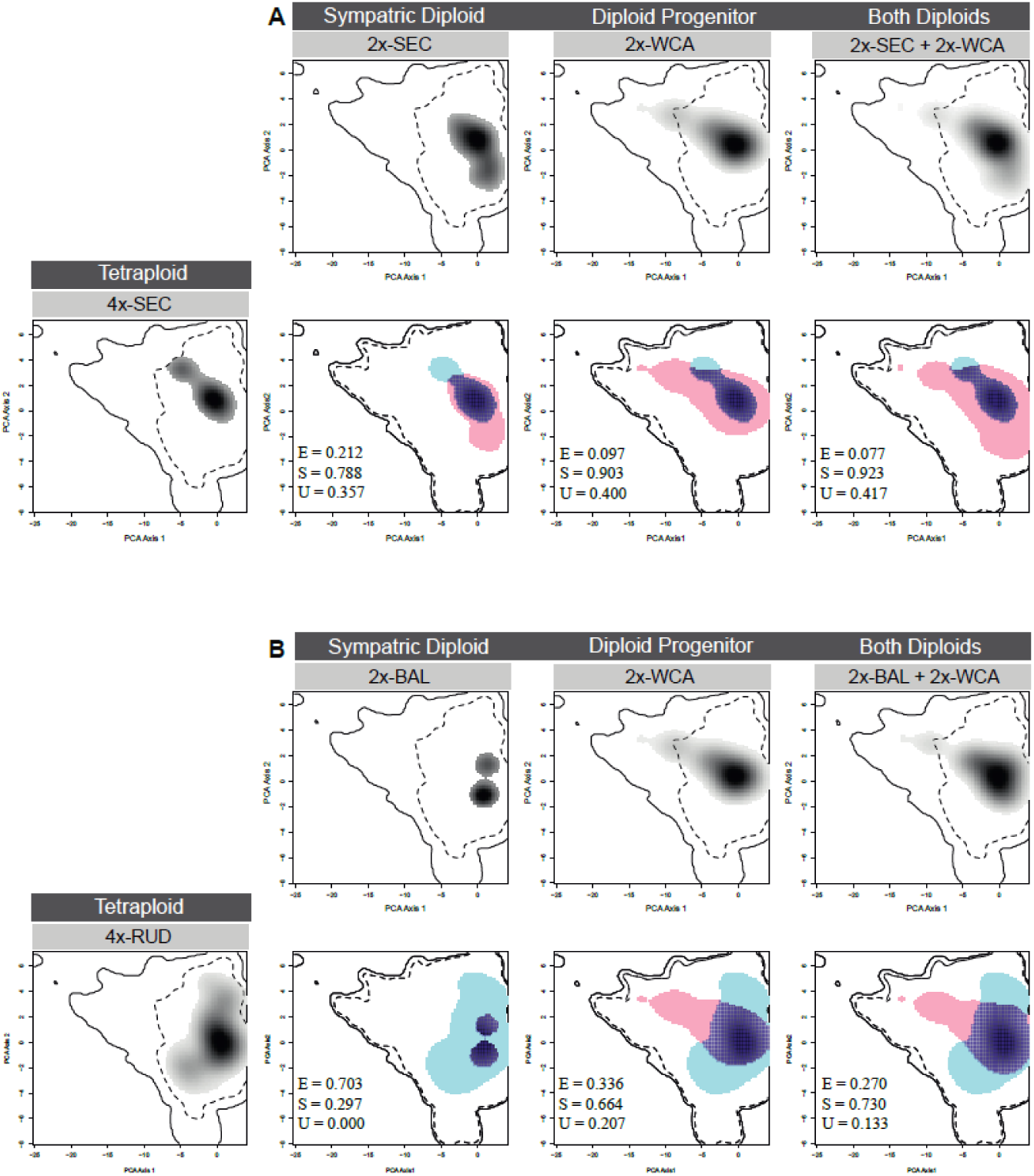
Climatic niche comparison of observed and expected niches for SE Carpathians tetraploid (**A**) and Ruderal tetraploid (**B**) lineages. Occurrence density grid for each diploid and tetraploid lineage is represented in the climatic space along the first two principal components (black & white plots). The observed niche of each tetraploid lineage is compared to the observed niche of their respective putative diploid ancestors and to their expected niches, which are calculated as the combined niche spaces of diploid ancestors. The fraction of the tetraploid climatic niche that overlaps with diploid ancestors indicates niche stability (S, dark blue), whereas the fraction that remains unfilled represents tetraploid niche contraction (U, pink). Climatic space occupied by the tetraploid lineages while not predicted by the combination of diploid progenitors indicates niche expansion (E, blue).

Comparing the observed niche of 4x-RUD to those predicted by the combination of niches of both diploid lineages that contributed to its genetic make-up (2x-WCA and 2x-BAL), we observe differences with respect to the previous comparison including just the ancestral diploid 2x-WCA lineage (Fig. 3m, Fig. 5b). We observe a lower niche expansion of 4x-RUD lineage with respect to both diploid lineages (E = 0.270). A lower unfilling index value (U = 0.133) and a higher stability (S = 0.730) are also shown (Fig. 5B).

## DISCUSSION

### 1. Varying extent of niche differentiation and expansion across tetraploid intraspecific lineages

It is usually assumed that polyploids have a different ecological niche than diploids, reflecting a WGD-driven shift in important functional traits, which in turn strengthens prezygotic isolation between ploidies and contributes to polyploid speciation (Levin, 2004; Husband et al., 2016). However, the existence and potentially the extent of ploidy-related niche differentiation differ between individual species and sometimes even among case studies (Glennon et al., 2014, Visger et al., 2016, Muñoz-Pajares et al., 2018, Castro et al., 2019). One explanation for this controversy is that niche evolution of polyploids might be detectable only when polyploid lineages are compared with their corresponding diploid ancestors but, unfortunately, the intraspecific genetic structure of polyploid complexes is often ignored when testing climatic niche evolution of polyploids. To test for this effect, we compared niches of ploidy cytotypes of *A. arenosa* - a species surrounded with controversy regarding the level of niche differentiation of its diploid and autotetraploid cytotype (Molina Henao & Hopkins, 2019, Morgan et al., 2020) - using the largest population sampling to date coupled with genotyping of each population. We assigned each autotetraploid population to one of the five major genetic lineages based on our genomic analyses (Fig. 2), which corresponded with previous genetic structuring results based on smaller sampling (Arnold et al., 2015; Monnahan et al., 2019) and put our results in the context of polyploid origin and interploidy gene flow by integrating with previous population genomic investigations involving both ploidies (Monnahan et al., 2019; Fig. 1).

In our study, niche shift was not supported when we compared diploid and tetraploid cytotypes globally, i.e. ignoring the intraspecific genetic structure (Fig. 3b, Table 1). However, we identified highly variable patterns of niche differentiation when the climatic niche of each tetraploid lineage was compared independently to the niche of the ancestral diploid lineage (2x-WCA). Niche conservatism between diploids and tetraploids was only found within the zone of primary coexistence (W Carpathians), where tetraploids of *A. arenosa* originated. This result corresponds with previous studies that found neither regional climatic nor local habitat differences between diploid and tetraploid populations of *A. arenosa* in this contact zone (Wos et al., 2019; Morgan et al., 2020). The fact that diploids and tetraploids do not show niche differentiation indicates that tetraploids can coexist in the same niche as their diploid progenitors. These results confirm growing body of empirical works showing that polyploids and diploids living in similar ecological niches tend to be common in autopolyploids (Godsoe et al., 2013; Glennon et al., 2014; Kirchheimer et al., 2016; Castro et al., 2019). Some such autopolyploids can escape or reduce competition with diploid progenitors by other changes in phenology, pollinators or parasite interactions (Thompson et al., 2014; Segraves & Thompson, 1999), or by post-pollination prezygotic barriers involving pollen competition and altered pollen-stigma interactions (Husband, 2016; Castro, Loureiro, Husband & Castro, 2020b). In the autotetraploid *A. arenosa*, other processes than niche differentiation might promote reproductive isolation between cytotypes, which in turn allowed tetraploids to get established in sympatry with their diploid progenitors. On the other hand, we cannot rule out the possibility of ongoing interploidy gene flow in this zone, which might mitigate niche differences between diploids and tetraploids. After the postulated single origin ∼20–30 thousands of generations ago (Monnahan et al., 2019), tetraploids of *A. arenosa* spread across Europe and diversified into four additional lineages (Fig. 1). When niches of these allopatric tetraploid lineages are compared to the ancestral diploid lineage (2x-WCA), the results largely vary among lineages (Fig. 3). While tetraploids from SE Carpathians (4x-SEC) exhibited a considerable niche contraction, the Alpine, Ruderal and C European tetraploid lineages have significantly expanded their niche with respect to the ancestral diploids (Table 1, Fig. 3). Furthermore, when we compared the climatic niches of these diverged tetraploid lineages to the tetraploid lineage that still occupies the ancestral area (4x-WCA) we also observed significant differentiation, suggesting notable post-WGD evolution of tetraploid niches (Table 1, Fig. 4). Most of them have experienced some niche expansion with respect to the ancestral 4x-WCA (Table 1, Fig. 4), and they have colonized very different climatic conditions either towards drier and warmer (4x-RUD and 4x-CEU), colder (4x-SEC), or more humid environments (4x-ALP).

Overall, our study confirms that niche evolution of polyploids is detectable only when polyploid lineages are compared with their corresponding diploid ancestor lineage, not globally. While niche similarity is observed when comparing the niches of cytotypes coexisting in the ancestral area of the tetraploid cytotype, niche expansion is mainly driven by post-WGD diversification into several lineages, primarily those that colonized warmer and drier anthropogenic habitats. These results demonstrate that niche shift is likely not driven by WGD *per se* in *A. arenosa* but rather reflects dynamic post-WGD evolution in the species, involving tetraploid migration and potential further interactions of tetraploids with other diploid lineages as is discussed in the next section.

### 2. Contribution of interploidy introgression to tetraploid niche expansion

Previous analyses demonstrated that genetic distinctness of particular lineages (4x-SEC, 4x-RUD) reflects strong interploidy gene flow after secondary contact with distantly related diploid lineages that have diverged before the WGD event (Monnahan et al., 2019). Such interploidy gene flow could have increased phenotypic and genetic variation which may allow tetraploids to colonize new ecological niches, in a way that is somewhat analogous to expectations stemming from hybrid origin of allopolyploids (e.g. Parisod & Broennimann, 2016; Huynh, Broennimann, Guisan, Felber & Parisod, 2020). Accordingly, the effect of introgression can be tested using an ecological niche comparison approach by looking at changes in terms of niche dynamic components such as unfilling and expansion. Here we leveraged the exceptionally well-described evolutionary history of *A. arenosa* involving localized post-WGD introgression from divergent diploids into two distinct tetraploid lineages (4x-SEC and 4x-RUD, Fig. 1; Monnahan et al., 2019) to assess if considering a niche of such additional diploid “donor” in addition to the 2x-WCA ancestor help to explain inter-ploidy niche divergence. Although notable effects have been observed in both cases, the results, once again, differed for each lineage. In the case of the 4x-SEC lineage, adding niche requirements of the sympatric 2x-SEC lineage do not explain tetraploid expansion but rather indicate tetraploid’s contraction in terms of temperature. These results show that interploidy gene flow between the 4x-SEC and the sympatric but evolutionary distant 2x-SEC has not led to a broader niche of the 4x-SEC lineage (Fig. 5a). On the other hand, introgression of the Baltic diploid lineage (2x-BAL) into the 4x-RUD lineage has slightly increased the ecological range of tetraploids towards colder conditions (Fig. 5b). This partly explains the massive expansion of the tetraploid Ruderal lineage towards more northerly habitats that are characteristic for the Baltic lineage. Indeed, previous studies have demonstrated that introgression of alleles involved in flowering time regulation from 2x-BAL to 4x-RUD has played a role in the early flowering of tetraploid ruderal *A. arenosa* plants (Baduel, Hunter, Yeola & Bomblies, 2018). Our results suggest a potential effect of interploidy gene flow in patterns of climatic niche evolution of tetraploid ruderal plants of *A. arenosa*. Thus, our study is one of the few examples supporting that interploidy introgression may be an important process for ecological adaptation of wild plants to challenging environments.

## Acknowledgements

We are particularly grateful to Jakub Vlček for his help and contribution to the genomic analyses. This work was funded by the Czech Science Foundation GAČR (19-06632S to KM), the Slovak Research and Development Agency (APVV, grant APVV-17-0616 to KM) and the OP RDE project CZ.02.2.69/0.0/0.0/18_053/0016976 (International mobility grant to NPG at Charles University). EZ, VZ, ML and FK were supported by long-term research development project RVO 67985939 (Czech Academy of Sciences). Computational resources were supplied by the project “e-Infrastruktura CZ” (e-INFRA CZ LM2018140) supported by the Ministry of Education, Youth and Sports of the Czech Republic. Access to CESNET storage facilities provided by the project “e-INFRA CZ” under the programme “Projects of Large Research, Development, and Innovations Infrastructures” LM2018140), is also acknowledged.

## Author contributions

NPG, GŠ, FK and KM conceived the ideas; GŠ, EZ, ML, FK and KM collected most of the plants; GŠ performed the lab work; VZ proceed the raw genomic data; MŠ wrote the script for pruning the input data for STRUCTURE and updated the script for detecting the convergence among replicates; JC wrote the script for calculating niche optimum and breadth; NPG analysed the data; NPG led the writing.

## SUPPLEMENTARY INFORMATION

**Figure S1.**
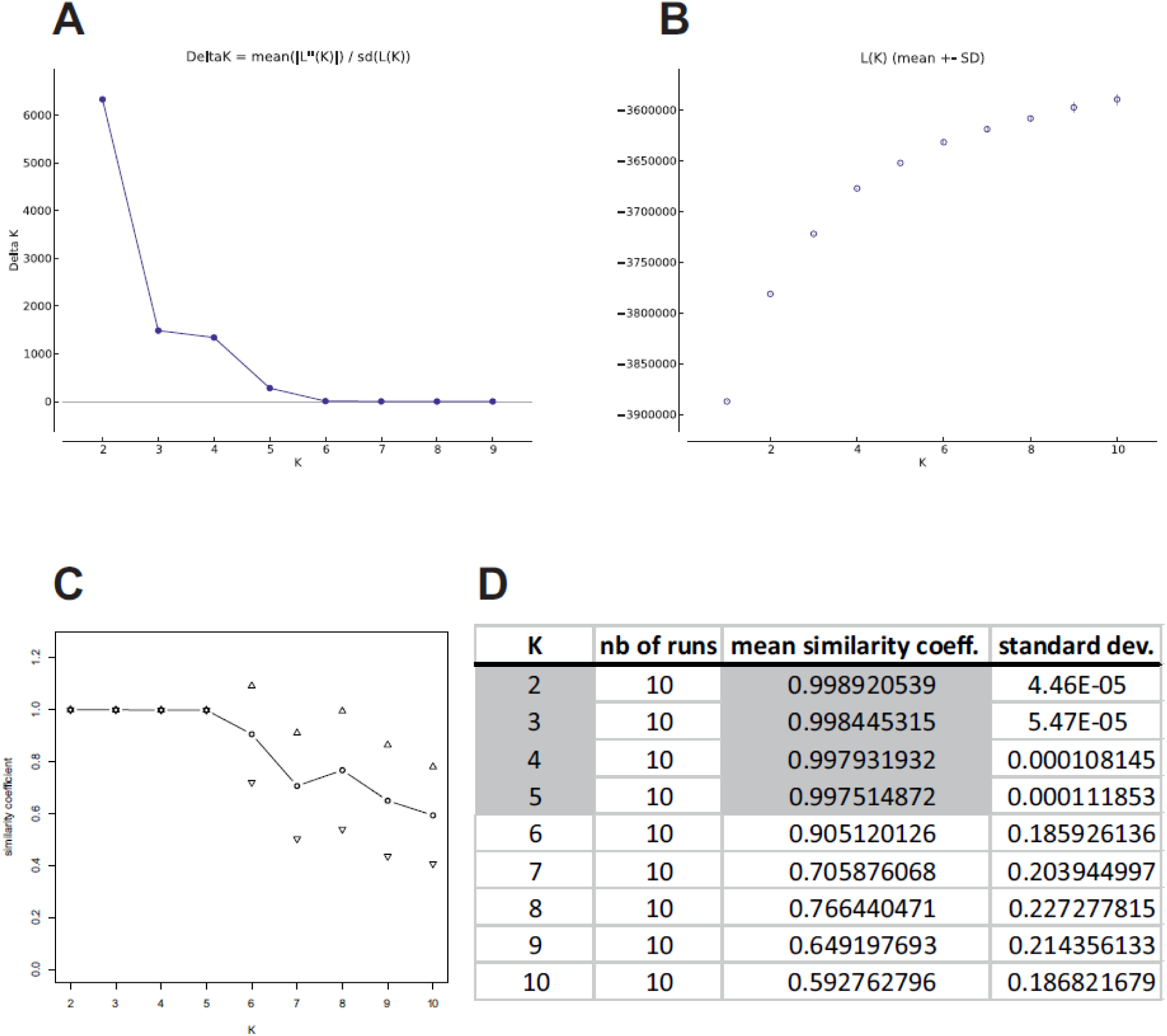
Summary of delta *K*, likelihood and convergence values of STRUCTURE replicates of tetraploid *A. arenosa* SNP dataset; **A, B** Mean delta *K* and likelihood values respectively, both calculated based on the Evanno method (Evanno, Regnaut & Goudet 2005), **C, D** Plot and values of mean similarity coefficients and standard deviation considering 10 replicates for each value of *K*.

**Figure S2.**
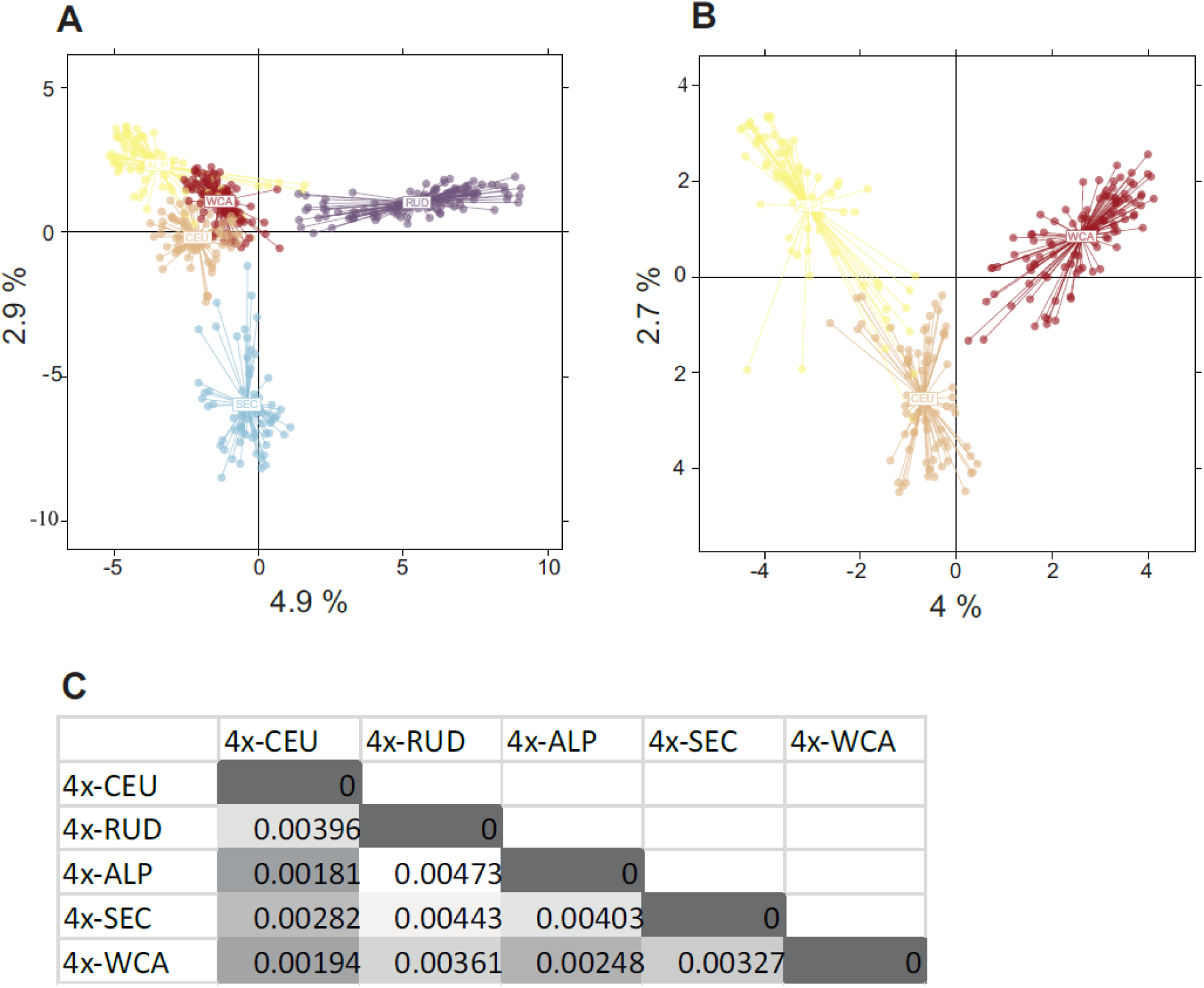
**A** Principal Components Analysis (PCA) of all tetraploid *A. arenosa* lineages identified in this study. Axis 1 and axis 2 explains 4.9% and 2.9% of the variation, respectively, **B** PCA including W Carpathians, C European and Alpine tetraploid lineages of *A. arenosa*. Two first axes explain 4% and 2.7% of the total variance, respectively. **C** Nei’s genetic distances among the identified lineages.

**Table S1**. Details of the *Arabidopsis arenosa* populations included in this study. Ploidy, lineage assignment, population code, number of individuals used for genomic analyses (*N*_ga_), populations used for niche comparison analyses (*N*_nc_) and locality details for each population are shown.

**Table S2**. Details on tetraploid *Arabidopsis arenosa* populations analyzed in this study. Population code, number of genotyped samples per population (sample size), ploidy level, methodological approach used to generate the data, lineage assignment and admixture proportions obtained by STRUCTURE are shown.

**Table S3**. Number of occurrences per cytotype and per lineage included in the niche quantification analyses before and after the filtering (closer than 10-km distance were removed).

**Table S4**. Description of environmental variables extracted from WorldClim and their contribution to the two first axes of the PCA-env.

